# SpecSeg: cross spectral power-based segmentation of neurons and neurites in chronic calcium imaging datasets

**DOI:** 10.1101/2020.10.20.345371

**Authors:** Leander de Kraker, Koen Seignette, Premnath Thamizharasu, Bastijn J. G. van den Boom, Ildefonso Ferreira Pica, Ingo Willuhn, Christiaan N. Levelt, Chris van der Togt

## Abstract

Imaging calcium signals in neurons of awake, behaving animals using single- or multi-photon microscopy facilitates the study of coding in large neural populations. Such experiments produce massive datasets requiring powerful methods to extract responses from hundreds of neurons. We present SpecSeg, a new open-source toolbox for 1) segmentation of regions of interest (ROIs) representing neuronal structures, 2) inspection and manual editing of ROIs, 3) neuropil correction and signal extraction and 4) matching of ROIs in sequential recordings. SpecSeg uses a novel method for ROI registration, based on temporal cross-correlations of low-frequency components derived by Fourier analysis, of each pixel with its neighbors. The approach is insightful and enables ROI detection around neurons or neurites. It works for single- (miniscope) and multi-photon microscopy data, eliminating the need for separate toolboxes. SpecSeg thus provides an efficient and user-friendly approach for analyzing calcium responses in neuronal structures imaged over prolonged periods of time.

## Introduction

The advances *in vivo* dual-photon fluorescence microscopy (Engert and Bonhoeffer, 1999; Svoboda et al., 1996) and the development of genetically encoded fluorescent biosensors (Chen et al., 2013; Mank et al., 2008) have revolutionized neurobiological research over the last two decades. The combination of these techniques has enabled the imaging of neuronal activity in awake behaving animals over timespans up to many months. This provides combined anatomical and functional information at the cellular level for hundreds of neurons at the same time, or at the dendritic, axonal or synaptic level in more restricted numbers of neurons (Cichon and Gan, 2015; Gambino et al., 2014; Iacaruso et al., 2017; Jaepel et al., 2017; Jia et al., 2010; Petreanu et al., 2009; Szalay et al., 2016; Wilson et al., 2016; Winnubst et al., 2015). More recently, gradient-index lens technology has made it possible to develop miniaturized single-photon fluorescence microscopes, or miniscopes. These are sufficiently light to be mounted on small, freely moving animals such as bats, birds and rodents, further expanding the possibilities for analyzing neural activity during behavior (Aharoni et al., 2019; Cai et al., 2016; de Groot et al., 2020; Ghosh et al., 2011; Liberti et al., 2017; Resendez et al., 2016). The insight these approaches provide about the functions and interactions of specific neuronal subtypes in different brain regions of awake behaving animals was previously unthinkable. The most-used approach is the imaging of changes in intracellular calcium levels as a proxy for neuronal activity. This is achieved by making use of genetically encoded calcium sensors such as GCaMP6 (Chen et al., 2013), whose fluorescent properties change upon binding calcium. The mere size of the obtained datasets, which consist of movies of calcium-indicator fluorescence images, forms a considerable challenge for data analysis. An important step in the analysis of calcium imaging data is the identification of cell bodies or neurites in the image sequences. Ideally, one can identify the same structures in recordings performed at different days over prolonged periods of time, enabling the assessment of changes in neuronal responses during learning at the single cell level. Identification of these regions of interest (ROI) is preferentially done in an automated fashion, as manual segmentation is neither reproducible nor scalable. Moreover, human annotators tend to include non-active ROIs and miss active ROIs with low background fluorescence (Giovannucci et al., 2019; Pachitariu et al., 2017). Automated ROI identification requires robust detection algorithms with minimal assumptions on the properties of ROIs to detect the circumferences of individual cells and neurites.

Various software packages have been published that accomplish this task, using different methods. Cell boundaries may be detected by multiple coupled active contours (Reynolds et al., 2017). Matrix factorization approaches (Giovannucci et al., 2019; Maruyama et al., 2014; Mukamel et al., 2009; Petersen et al., 2018; Pnevmatikakis et al., 2016; Zhou et al., 2018) determine the activity of neurons and their delineation by considering fluorescence as a spatiotemporal pattern that can be expressed as the product of a matrix encoding location and a matrix encoding time. Deep learning approaches (Apthorpe et al., 2016; Klibisz et al., 2017) define ROIs based on neuron features learned from data in which cells were manually detected. Dictionary learning approaches make use of neuron templates to identify ROIs (Pachitariu et al., 2017). Finally, correlation-based approaches define ROIs based on activity correlations between pixels (Kaifosh et al., 2014; Smith and Häusser, 2010; Spaen et al., 2019). All these methods have their strengths but also some weaknesses. For example, deep learning approaches that select ROIs based on shapes must be trained for different types of data and may include neurons from which no signal can be extracted. Identifying ROIs with highly variable shapes is complicated for most approaches except those based on activity correlation. For most methods, it is necessary to introduce significant adaptations to the software in order to make it suitable for specific experimental settings and the underlying calculations that define the ROIs can be difficult or impossible to track, making it difficult to select the optimal settings. Correlation-based approaches are easy to understand and require few predefined constraints but can be severely limited by noise in the recordings.

In this paper, we describe an open-source pipeline (Fig.1) for calcium imaging data analysis. It makes use of a novel approach for ROI detection, termed SpecSeg, which is based on cross-correlations of low frequency components, derived through Fourier transforms, of each pixel trace with its eight adjacent neighbors. It makes use of our finding that low-frequency fluctuations below 0.4 Hz are a signature of active neurons. Our approach is insensitive to noise, straightforward and highly insightful for the end user as it enables the visualization of the ROI detection process. It enables the detection of ROIs around irregular structures such as dendrites and axons and the parameters set by the users are intuitive. The pipeline also includes a user interface for quality control and the manual splitting, addition or rejection of ROIs, and a tool to match ROIs in sequential imaging sessions (Fig. 1). Finally, the pipeline can be used for the analysis of data acquired using multi-photon microscopes and single-photon miniscopes. Together, this new pipeline provides an efficient and user-friendly approach for analyzing calcium responses in neuronal structures imaged over prolonged periods of time.

**Figure 1.**
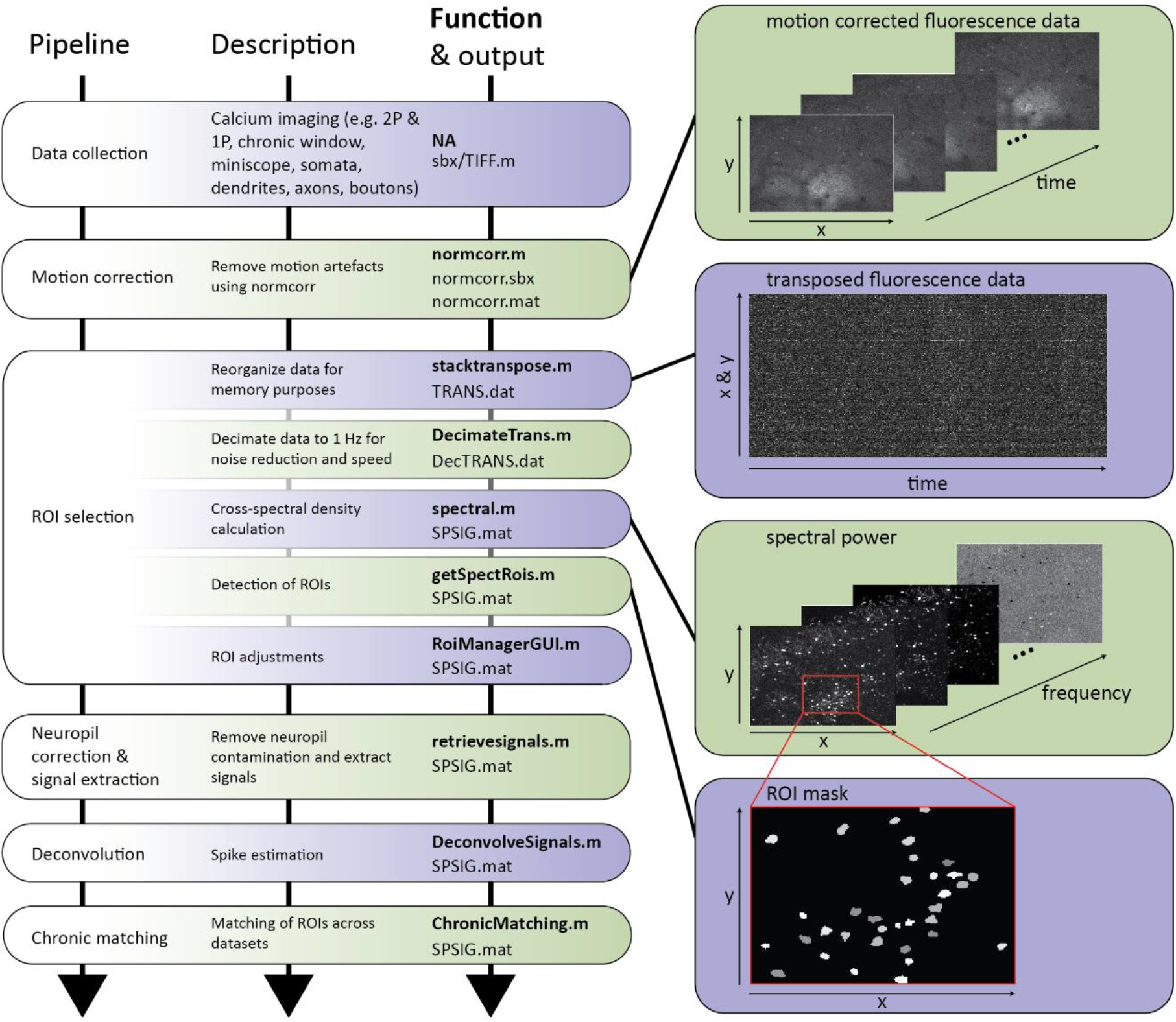
Overview of the pipeline for automated region of interest (ROI) selection and signal extraction. After collection, calcium imaging data needs to be motion corrected using NoRMCorre. ROI selection involves data reorganization, extraction of frequency components of pixel traces and the drawing and refinement of ROIs around the peaks identified in the images based on the frequency components. A user interface enables the manual rejection, splitting or adding of ROIs. Next, neuropil is subtracted and the signal extracted and deconvolved. A toolbox for matching of ROIs in repeated recordings is also provided.

## Results

### Pipeline

Figure 1 shows the components of our pipeline for the analysis of chronic calcium imaging datasets. The pipeline requires motion-corrected calcium imaging data in raw TIFF or SBX (Neurolabware) format. We use an adapted version of NoRMCorre (Pnevmatikakis and Giovannucci, 2017) to motion correct our SBX files. A motion-corrected series of TIFF images can also be used, but this first needs to be converted to SBX format for which we provide a conversion tool.

The first step in the pipeline is ROI selection, which involves (a) the reorganization and downsampling of the data in order to speed up memory retrieval, (b) extraction of the frequency components of the fluorescence traces for each pixel (“pixel trace”) by Fourier analysis and the creation of images representing how well they correlate with those of neighboring pixels (c) the identification of peaks within the images and the construction of preliminary contours (ROIs), and (d) further refinement of the ROIs based on activity correlations within each contour. The second step in the pipeline is the ROI manager, a user interface for the inspection of the ROIs that has tools for manually rejecting, splitting or adding ROIs. The third step is neuropil subtraction and signal extraction and the final, fourth step is deconvolving the signals, using maximum likelihood spike estimation (MLSpike, Deneux et al., 2016).

In addition, a separate toolbox is included for matching the selected ROIs in sequential imaging sessions by aligning the images and measuring the overlap of ROIs between different sessions. The sensitivity of ROI matching can be changed easily with an overlap threshold and the results of the matching can be evaluated and edited. Below, each step in the pipeline is described in detail. The MATLAB implementation of the toolbox and instructions on how to install and use the software can be found at https://github.com/Leveltlab/SpectralSegmentation.

### ROI selection based on cross-spectral power

We established a novel approach for the automated segmentation of ROIs in calcium imaging data based on cross-spectral power of the pixel trace. To develop the method, we made use of calcium imaging data acquired repeatedly using two-photon microscopy in mice expressing the genetically encoded calcium indicator GCaMP6f in the primary visual cortex (V1) and tested it on a variety of different datasets from different brain regions and using different imaging approaches.

The first step in the process (stacktranspose.m) is to transpose the image sequences to place time in the first dimension and width * height in the second dimension, and to down-sample the data to approximately 1Hz (DecimateTrans.m), using the MATLAB decimate function (Mathworks^®^). This way, each pixel trace is organized in sequential order in memory and can be accessed rapidly, speeding up the next steps in the process.

Next is the cross-spectral power calculation (spectral.m). Each pixel trace is cut into overlapping one-minute segments, and a discrete Fourier transformation is applied to each segment to extract frequency components between 0.013 and 0.5Hz, with a bin width of 0.017Hz. Then we calculate the cross-spectral density function of each pixel with its 8 neighbors, and average these over all segments. Finally, the average cross-spectral density functions are normalized with the variances of each pixel and its neighbors, and the average cross-spectral power for each frequency component at each pixel is calculated (this estimate is a measure for how well pixels correlate with their direct neighbors at each frequency component). This results in a series of 30 images, each representing a different frequency component.

These images of spectral components provide a better basis for the selection of ROIs around active neurons than the fluorescent signal (Fig. 2A-B). We find that spectral components below 0.4Hz are the most indicative for active neural elements (Fig. 2C-I, Fig. 3A-C). As a consequence, when we compose images by translating the cross-spectral power of a particular low-frequency component into brightness, active elements in the image become very clearly separated from the background (Fig. 2C-F), even when their level of fluorescence is low (see neuron in red box). Interestingly, active neurons also look different in cross-spectral images and do no longer have a dark central nuclear region. Low-frequency pixel correlations tend to show a central maximum that declines to the border of a neuron (Figs. 2, 3). In contrast, non-active neurons with high levels of fluorescence, which are visible in the average fluorescence projection (see neuron in cyan box, Fig. 2A, J), do not show up in the spectral images. ROIs can thus be selected by searching for the largest local maxima in these spectral images and drawing contours around them (Fig. 3). Because the images of the different frequency components reveal different neurons and/or neuronal compartments, a complete set of ROIs is created by adding up all ROIs detected in the spectral images of all low-frequency components (Fig. 2G, H).

**Figure 2.**
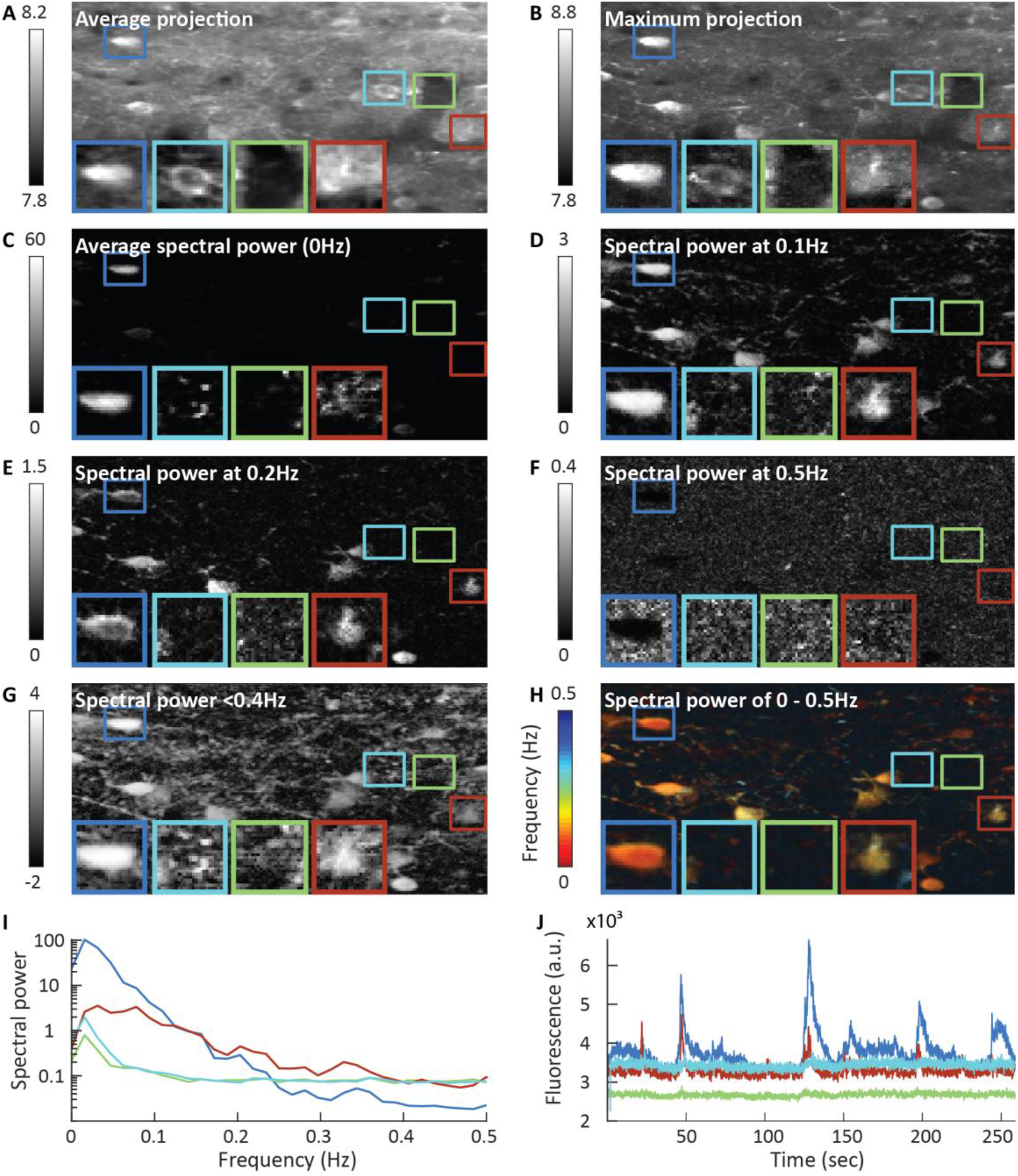
Comparison of fluorescence and spectral power images. **A**) Average fluorescence projection. The colored squares indicate regions containing an easily recognizable active neuron (dark blue); a silent neuron (cyan); no neuron (green); a hardly recognizable but active neuron (red). **B**) Maximum fluorescence projection. Note that the neuron in the cyan region can be easily detected, while the neuron in the red region is hardly visible **C-H)** Cross-spectral images. Note that the silent neuron in the cyan region is hardly visible in the cross-spectral images, while the active cell in the red region can be easily identified **I)** The cross spectral power of the four example areas. Note the logarithmic y-axis. **J)** Pixel traces of the example areas denoted in panel A-H.

**Figure 3.**
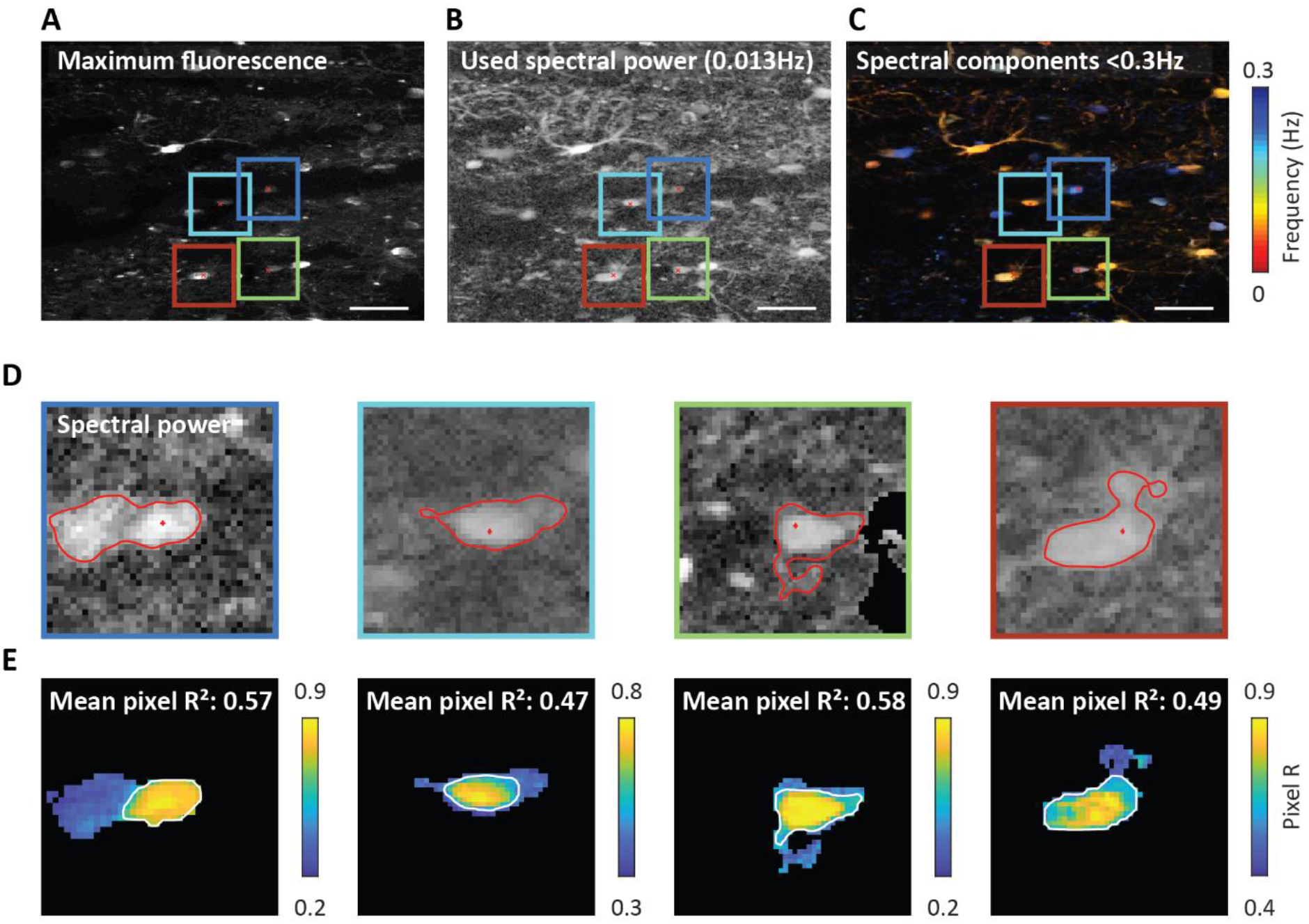
Automatic ROI creation and refinement. **A)** Maximum-fluorescence projection of the recording. The colored squares denote four example areas that are selected for an ROI search based on local maxima of the cross-spectral image in the next panels. Scale bar = 40µm. **B)** The spectral power of the first frequency component (0.013 Hz) that is used in the ROI search. **C)** Cross-spectral power, with different frequency components displayed in different colors. **D)** The spectral power from panel B in the selected areas containing local maxima. The contour that is found around these putative neurons is displayed in red. For these neurons the threshold for pixel inclusion was too low, resulting in ROIs that are too large. These ROIs are automatically refined based on pixel correlation. **E)** The pixel correlations of the ROIs displayed in D. The color displays the pixel correlation of the signal from each pixel with the signal from the ROI local maximum. The pixel correlation threshold facilitates ROI refinement. The new contour, shown in white, is determined by the pixel correlation values.

### Automated ROI refinement

Some contours selected this way may still contain overlapping or closely juxtaposed neuronal elements (Fig. 3A-D), especially if the density of cell bodies or neurites is high. To prevent this problem, contours are further constrained based on whether the fluorescence pixel traces within the contour are tightly correlated as would be expected if they are from the same neuron. We do this by taking the median fluorescence trace of the pixel traces from the local maximum and its eight neighboring pixels and correlating this trace with each pixel trace in the preselected contour. This results in correlation values for every pixel in the ROI (pixel R). Next, using a threshold set at half the maximum correlation strength (this threshold can be adapted), a new contour is selected around the best-correlated pixels (Fig. 3E). To save an estimate of how well the fluorescence pixel traces are correlated inside an ROI, we square the pixel correlation values of an ROI and average them, resulting in the mean pixel R^2^ (Fig. 3E). The mean pixel R^2^ indicates how much of the pixel variance is explained by a shared signal within an ROI. We assume that this signal is the actual activity trace of a neuron, possibly including a general neuropil signal. The remaining variance is due to sources surrounding a neuron influencing individual pixel traces separately.

### Versatility of ROI selection approach

Restrictions can be applied in order to select ROIs around cell bodies, such as the minimal and maximal surface area of the ROI or its roundedness. Without these restrictions, thin elongated contours can be easily selected, making it also possible to define ROIs on dendrites and axons (Fig. 4A-C). We tested the ROI selection approach on various datasets, including two-photon calcium imaging using a GRIN-lens in visual thalamus (Fig. 4D) as well as single-photon miniscope imaging in pre-motor cortex (M2) and striatum (Fig. 4E-F). Single-photon miniscope signals are generally more correlated and contaminated by out-of-focus signals. Therefore, we included an additional background correction tool for single-photon imaging data (BackgroundSubtractSbx.m) (Fig. 4E-F).

**Figure 4.**
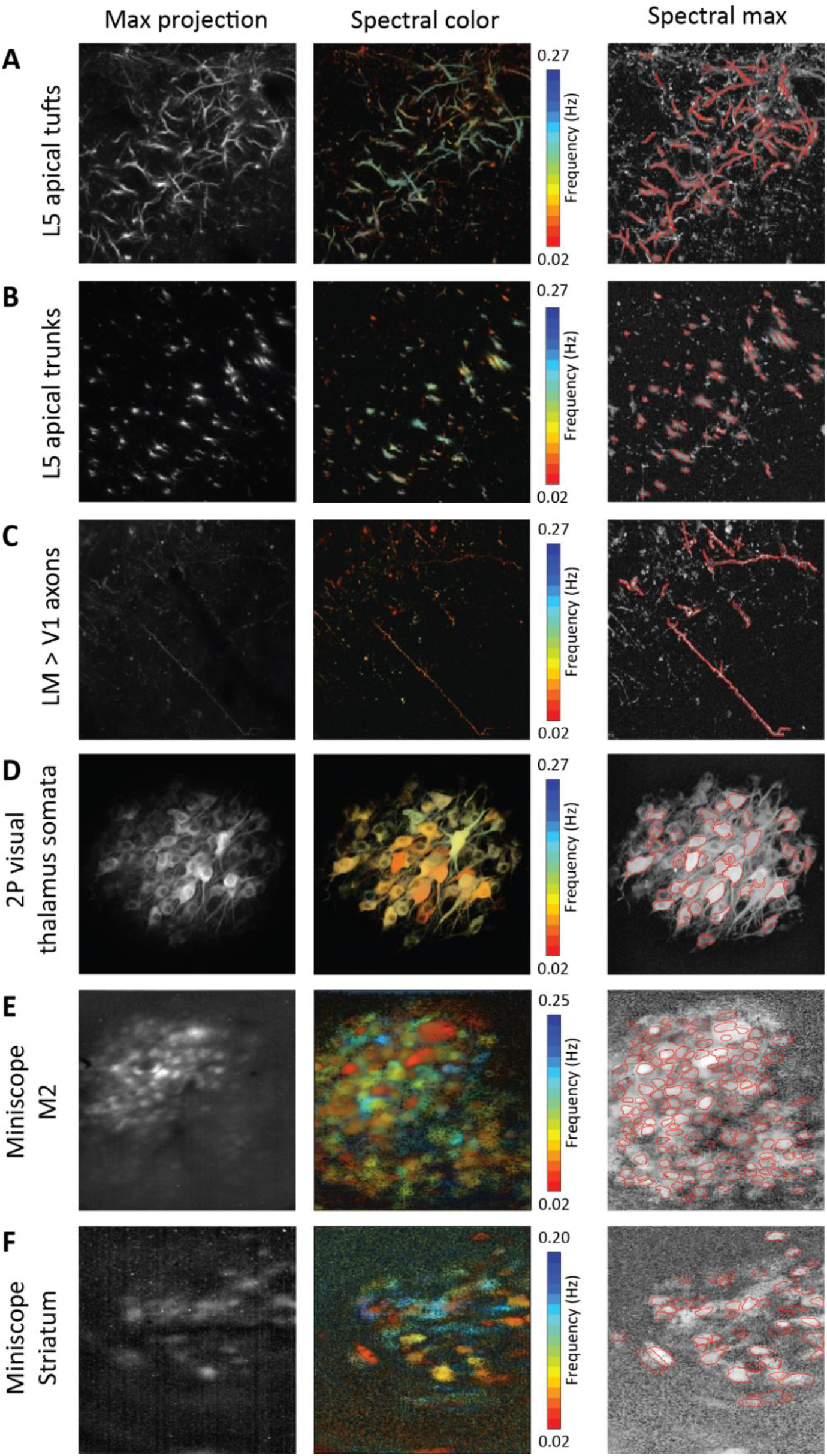
Examples of fluorescent signals, cross-spectral power and ROI selection in datasets derived from different brain regions and/or imaging techniques. Left column: maximum fluorescence projections. Middle column: cross-spectral power, with different frequency components in different colors. Right column: cross-spectral power maximum projection, with contours of ROIs in red. **A)** Two-photon microscopy of **l**ayer 5 apical tufts in mouse V1. **B)** Two-photon microscopy of layer 5 apical trunks in mouse V1. **C)** Two-photon microscopy of visual lateromedial area axon projections to V1. **D)** Two-photon microscopy of visual thalamus using a GRIN-lens. **E)** Single-photon miniscope imaging in mouse premotor cortex (M2). **F)** Single-photon miniscope imaging in mouse striatum.

### Comparing SpecSeg ROI selection in two-photon imaging data with other packages using Neurofinder

To compare the efficiency of our method with that of other packages that are available, we made use of Neurofinder datasets (http://neurofinder.codeneuro.org/). Our results (SegSpect0, by Spectral Segmentation) scored in the mid-range of all tested methods. Because this seemed a relatively low score considering the good results we obtained with SpecSeg on our own datasets, we looked at the Neurofinder results in more detail. We noticed that in several Neurofinder datasets, SpecSeg missed more than 50% of the Neurofinder “ground-truth” ROIs and found additional ROIs that were not included as ground-truth ROIs, both contributing to the lower score (Fig. 5A, D). To understand why SpecSeg selected different ROIs than those considered as ground truth, we calculated the mean pixel R^2^ of the pixels within the latter ROIs (Fig. 5B, C, E, F) (see methods). This provides an estimate of how much of the pixel variance is derived from a shared signal within an ROI. When we sorted the ROIs based on the mean pixel R^2^, we noticed a sharp decline of this estimate in the ground-truth ROIs in all of the Neurofinder datasets (Fig. 5C, F). In the Neurofinder datasets, most ROIs had a mean pixel R^2^ below 0.15, suggesting that > 85% of the signal variance in these ROIs originated from external sources or simply noise (Fig. 5G-H). ROIs selected by SpecSeg all had mean pixel R^2^ values well above this level. This resulted in only part of the ROIs defined as ground truth in the Neurofinder datasets to correspond with those drawn using SpecSeg. Importantly, SpecSeg missed less than 1% of the ground-truth ROIs with a mean pixel R^2^ over 0.3. SegSpec also selected additional ROIs representing active neurons that were not identified among the Neurofinder ground-truth ROIs. These ROIs had high pixel correlations and showed clearly defined responses with high signal to noise ratios (Fig. 5A, D). This shows that it is crucial to select ROIs based on activity and not solely on cell morphology visualized by the baseline fluorescence of the calcium indicator, as some responsive neurons may be invisible especially when using calcium indicators with very low calcium-free fluorescence such as JCaMP7c (Dana et al., 2019). In addition, we also compared our ROIs with those obtained using Suite2P (Pachitariu et al., 2017). The authors of Suite2P define a model that combines the estimated signal of an ROI with its neuropil background. Using a cost function to minimize the difference between model and actual fluorescence time course, the parameters of this model for each pixel are estimated. Again, we found overlapping subsets for ROIs with high mean pixel R^2^ values. The ROIs that SpecSeg did not select generally had a mean fluorescence trace similar to the background fluorescence in the Suite2P visualization tool.

**Figure 5.**
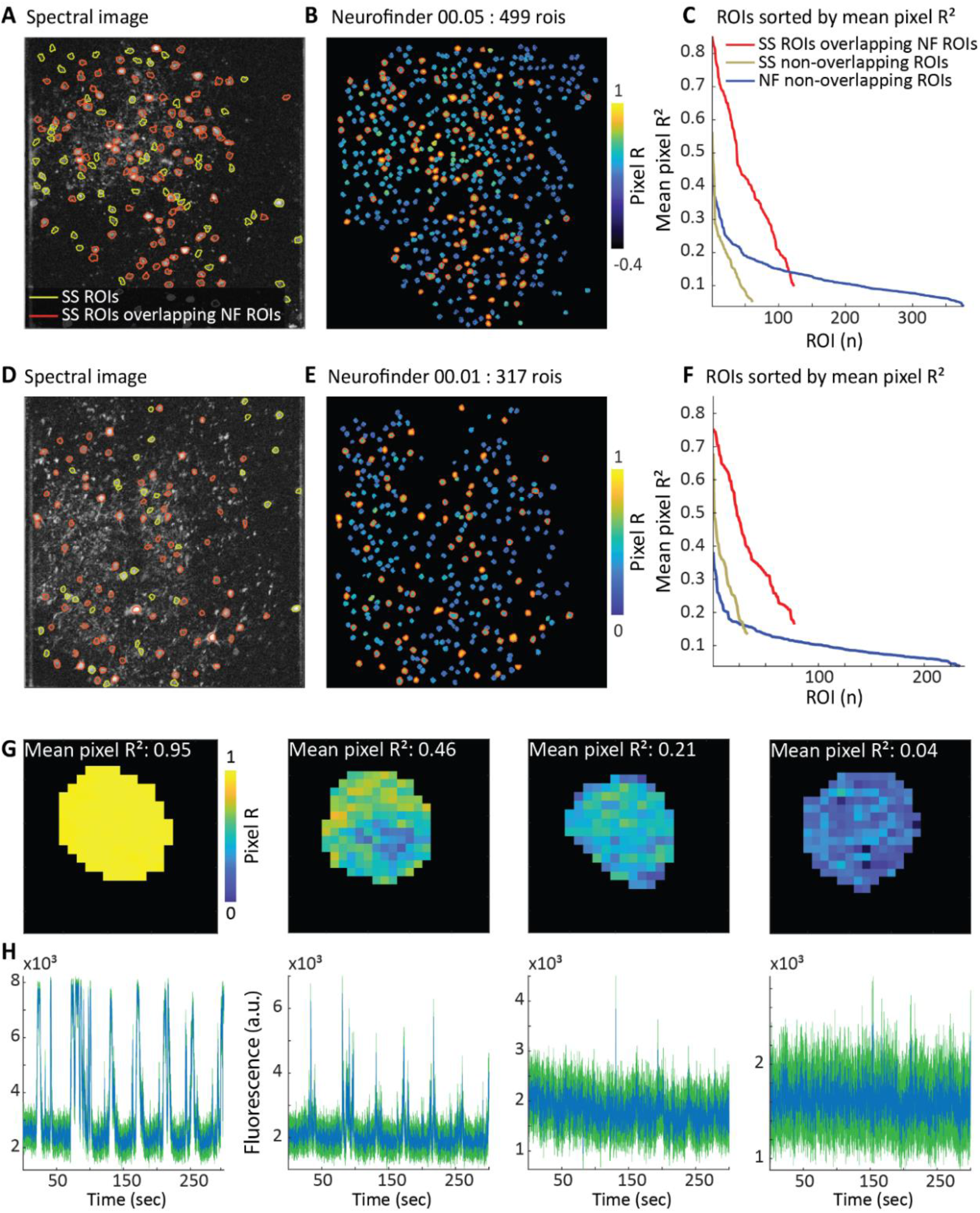
In Neurofinder datasets, SpecSeg only identifies ROIs from which a reliable signal can be extracted. **A)** Spectral image of a Neurofinder (NF) dataset, with ROIs encircled by red (overlapping) and green (non-overlapping) contours. **B)** Pixel R for ground truth ROIs for the same dataset. Color bar indicates pixel R. Red encircled Neurofinder ground truth ROIs overlap with SpecSeg (SS) ROIs. Note that SpecSeg selected all ROIs with the highest mean pixel R^2^. **C)** Mean pixel R^2^ of ROIs sorted by magnitude. Neurofinder ROIs overlapping with SpecSeg ROIs are plotted in red. The golden line displays SpecSeg ROIs that do not overlap with Neurofinder ROIs, while blue line represents Neurofinder ROIs that do not overlap with SpecSeg ROIs. **D, E, F)** Like A, B, C but for a different dataset. **G)** Correlation of each pixel trace within an ROI with the median pixel trace at the local maximum of cross-spectral power and its 8 neighboring pixels, of four example ROIs with decreasing mean pixel R^2^ from the 03.00 Neurofinder dataset. **H)** Fluorescence traces (blue) plotted against their SEM * 3 (green) for the four example ROIs. Note that around a mean pixel R^2^ of 0.2 or lower, the signal becomes non-significant.

Together, we suspect that ground-truth ROIs in the Neurofinder dataset are largely based on anatomical characteristics of GCaMP6-expressing neurons and include many silent neurons. Hence, the best performing methods on the Neurofinder datasets may use convolutional neural networks trained on ground truths that include neurons with low pixel correlations. Whether neurons that show so little activity that they cannot be reliably separated from noise or neuropil activity should be included in the analysis is questionable, though.

### Single-photon miniscope data analysis comparison with CNMF-E

Single-photon miniscope imaging data present additional problems for ROI selection and signal extraction. Single-photon data suffer heavily from out-of-focus fluorescence. The background fluorescence can contain fluctuations that are stronger than the signal from individual neurons themselves. The images also contain vignetting, meaning they are darker toward the edges. These problems require additional processing. A widely used single-photon analysis is CNMF-E (Zhou et al., 2018). With this method, ROIs are initialized based on a heavily filtered image, which is not suited for signal extraction. Therefore, the ROIs are refined and the signal estimated by constrained matrix factorization. SpecSeg uses a different approach, and first removes the out of focus fluorescence and vignetting from the video data itself via background subtraction. The background subtraction estimates background fluorescence for every frame individually, by filtering each frame with a large disk (∼200 µm). The resulting background fluorescence image is subtracted from the original frame (see methods). This approach decreases both out of focus fluorescence and vignetting, while small signal sources remain present in the data. The data from Fig. 4E-F was analyzed with both CNMF-E and SpecSeg. We found that 60% of the ROIs had spatial footprints that were more than 30% overlapping between the CNMF-E and SpecSeg analysis. Matched ROIs are shown in Figure 6A and E. Unlike in the two-photon Neurofinder dataset comparison, the mean pixel R^2^ was not significantly higher in the SpecSeg ROIs (Fig. 6B, F), probably because CNMF-E also selects ROIs based on activity, and not predominantly on anatomical features. The extracted signal traces from ROIs correlated well with the signal traces from corresponding ROIs from CNMF-E after background correction. Without background correction the signal traces correlate little with CNMF-E or the SpecSeg corrected signals (Fig. 6C, D, G, H).

**Figure 6.**
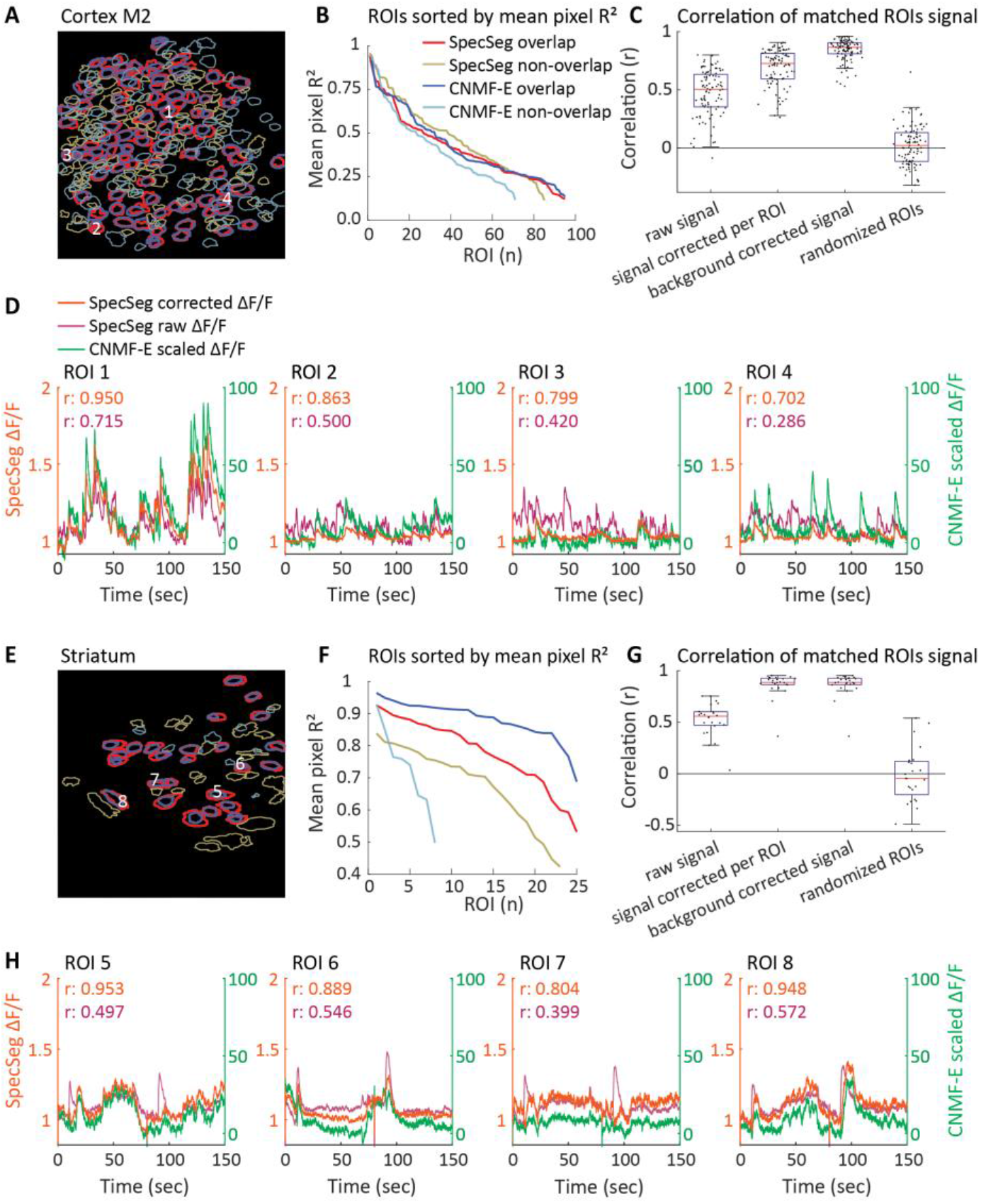
Comparison of CNMF-E and SpecSeg for the analysis of single-photon miniscope data. **A)** Miniscope calcium-imaging data from premotor cortex (M2) is analyzed using both SpecSeg and CNMF-E. ROIs identified by SpecSeg and CNMF-E are shown. Colors indicate whether or not the ROIs identified by the two methods overlapped (see panel B for color coding). White numbers, identify the overlapping ROIs whose signals are shown in panel D. **B)** Mean pixel R^2^ of ROIs sorted by magnitude. Overlapping CNMF-E and SpecSeg ROIs represent the same neurons. However, their spatial footprints are slightly different, also resulting in slightly different mean pixel R^2^. Non-overlapping ROIs do not have a significantly lower mean pixel R^2^ than overlapping ROIs. **C)** Signals extracted by SpecSeg are strongly correlated with the signals extracted by CNMF-E for overlapping ROIs. In contrast, correlating the raw signal with the CNMF-E signal results in lower r values. Background correction of the ROI signal with a surrounding donut ROI, as used for two-photon data, increases the correlation. Pixel-& frame-wise background correction increases the correlation even more, illustrating the importance of background correction. The fourth condition ‘randomized ROIs’ is a control and correlates the background corrected signal from the SpecSeg ROIs with random ROIs from CNMF-E. **D)** Signals from four example neurons, marked in panel A, are shown. The r values represent the spearman correlations between the ROIs’ SpecSeg signals and the CNMF-E signals. The correlation between the CNMF-E signal and the background-corrected SpecSeg signal (background-corrected signal in panel C) is indicated in orange, while correlation with the raw SpecSeg signal is indicated in magenta. **E)** Same as panel A but for calcium imaging data from striatum. **F)** Mean pixel R^2^ of ROIs sorted by magnitude. **G)** For the striatum data, ROI-wise signal correction is more effective than in the M2 data, because the striatum has a sparser population of neurons. This allows for more data to be used in the subtraction, making it more similar to the pixel-wise background correction. **H)** The signal from the four marked ROIs marked in panel E.

### Speed

In order to get an impression of the speed of our approach, we automatically timed the different components of the pipeline on two different computers analyzing 16 different datasets, varying in size from 6 to 51 gigabytes. Depending on the hardware that was used and the size of the dataset, the different components of the analysis pipeline varied in the time they required for completion (Fig. 7). This makes it difficult to provide hard numbers about the speed at which the SpecSeg pipeline processes the data. However, a useful indicator is that 1 hour of acquired data imaged at 15 Hz was processed by automated analysis in approximately 5 hours, from NoRMCorre motion correction until signal extraction. More than half of the time was used by the NoRMCorre motion correction. This illustrates that the speed of the SpecSeg pipeline is not a bottleneck, and in most cases performs the analysis overnight.

**Figure 7.**
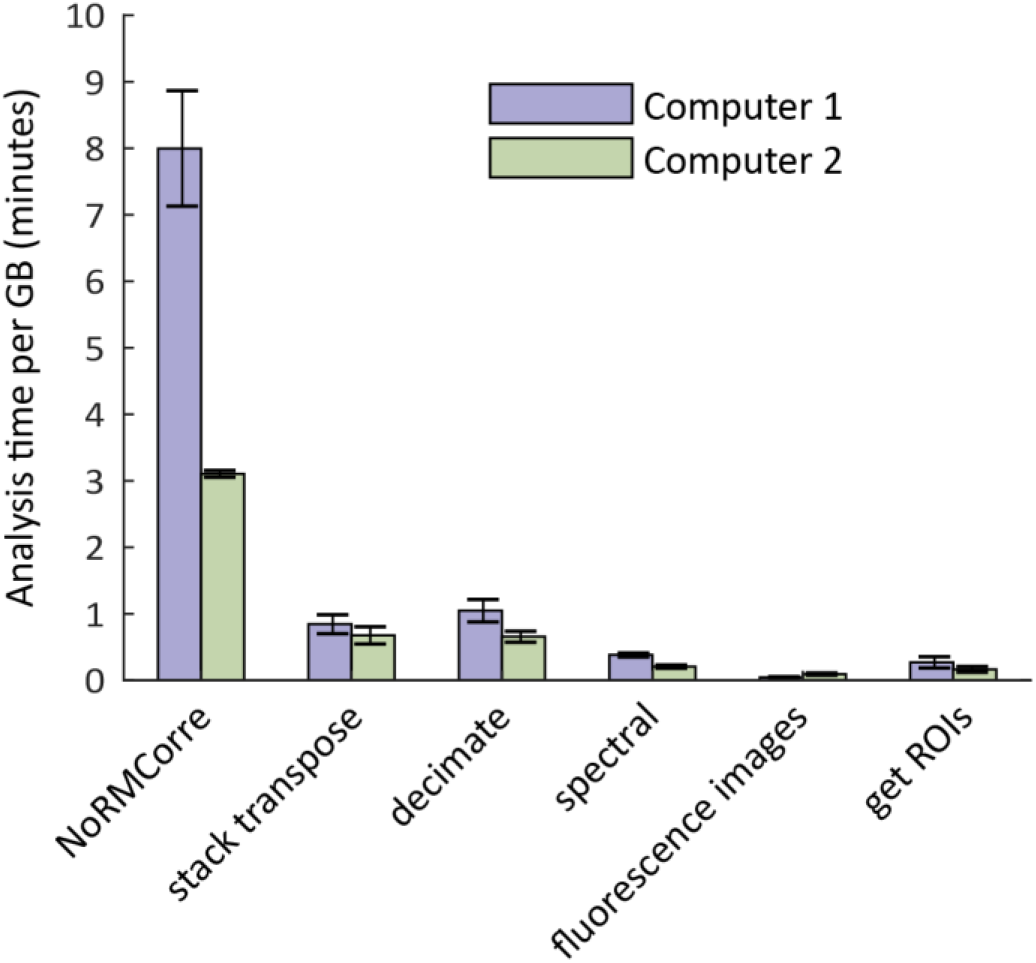
Timing analysis pipeline. Timing the analysis per GB of raw data with two different computer systems. Computer 1: 2x CPU: Intel Xeon E5-2620 2.4GHz, 6 cores. RAM: 96GB. Computer 2: CPU: Intel Core i7-4770 3.4GHz quadcore. RAM: 32GB. Total automated analysis time including motion correction with NoRMCorre. Computer 1: 10.6 minutes per GB (n=8). Computer 2: 4.5 minutes per GB (n=8). Not including NoRMCorre. Computer 1: 2.5 minutes per GB (n=8). Computer 2: 1.8 minutes per GB (n=8). Error bars shows the standard error of the mean.

### User interface for rejecting, splitting or manually defining ROIs

The automated procedure for selecting ROIs is highly effective. However, in some cases, ROIs can still be found that contain signals from multiple neurons or do not seem to represent a neuronal structure. Because such errors can cause misinterpretation of calcium imaging data, we developed a graphical user interface for splitting or deleting such ROIs. Additionally, because the automated ROI selection process is activity dependent, ROIs are not drawn around neurons that are not active in the imaging session. The user interface also makes it possible to add ROIs for such neurons (Fig. 8A), in case the experimenter wants to include them in their analysis – for example in a chronic experiment in which the neurons do show activity in other recording sessions.

**Figure 8.**
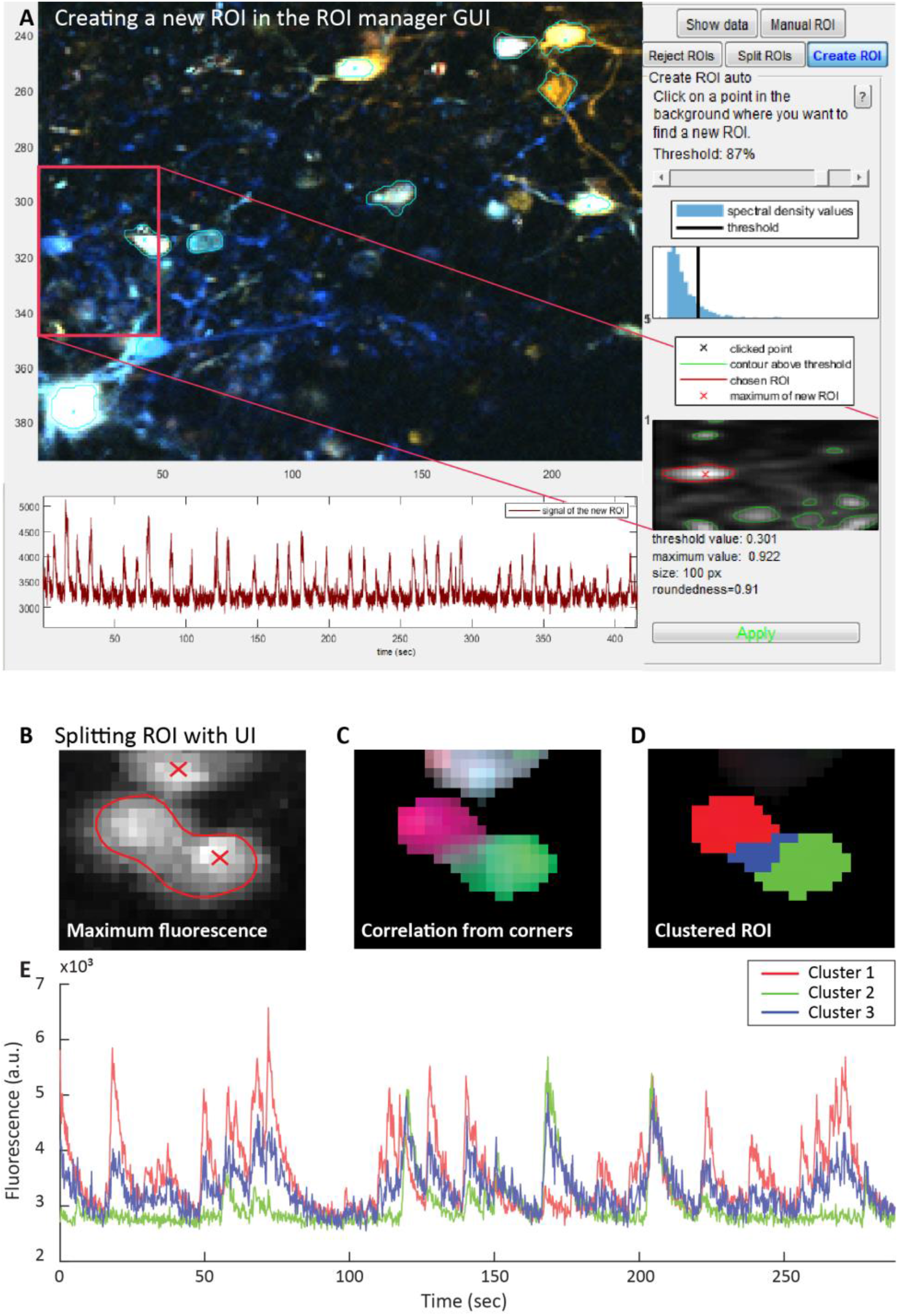
ROI manager user interface. **A)** A screenshot of the user interface. The functionality is set to ROI creation. The ROI creation function of the UI allows the user to add additional ROIs based on the current image in the UI. The current image shown in A is the cross-spectral power, in which several frequencies are shown in different colors. **B)** Maximum fluorescence of an ROI that envelopes two neurons. **C)** To guide and calculate the ROI splitting the fluorescence signal from the four corners is correlated with the all fluorescence pixel traces in the ROI. This results in 4 correlation values per pixel, which are shown in different colors. **D)** The ROI is clustered into three clusters with k-means clustering using the correlation values from **C**. The number of clusters is set by the user. **E)** The fluorescence signal of the three clusters.

Sometimes ROIs are created that envelop multiple neurons (Fig. 8B). To visually guide the splitting of these ROIs, pixel traces within the ROI are correlated with four reference points and correlation values are color coded. In case the ROI contains signals from more than one neuron, this becomes clearly visible (Fig. 8C). The correlation values are then used in k-means clustering to split the ROI into multiple ROIs, or to delete parts of the ROI, on the user’s request (Fig. 8D). ROIs can also be deleted entirely or marked to be kept when more stringent criteria for ROI selection are applied. Finally, ROIs can be drawn in manually, or added by the computer, based on the fluorescence or spectral image (Fig. 8A). Altogether, these additional tools provide the means to adjust the ROI selection process in an easy and user-friendly way.

### Neuropil correction and signal extraction

Once the ROIs are selected, the calcium signals from the cells can be extracted by averaging all pixel traces within the ROI contours. However, because the calcium signals from the neuropil can contaminate those of the cells, neuropil signal is first subtracted from the ROIs (Zhou et al., 2018). To achieve this, we select an area surrounding each ROI, taking care to avoid other ROIs (Fig. 9, see methods for details). The average of the pixel traces in this area is used as an estimate of the local neuropil activity. After multiplying it by 0.7 it is subtracted from the averaged fluorescence trace from the selected ROI. This method of neuropil subtraction is widely used (Chen et al., 2013; Khan et al., 2018; Tegtmeier et al., 2018).

**Figure 9.**
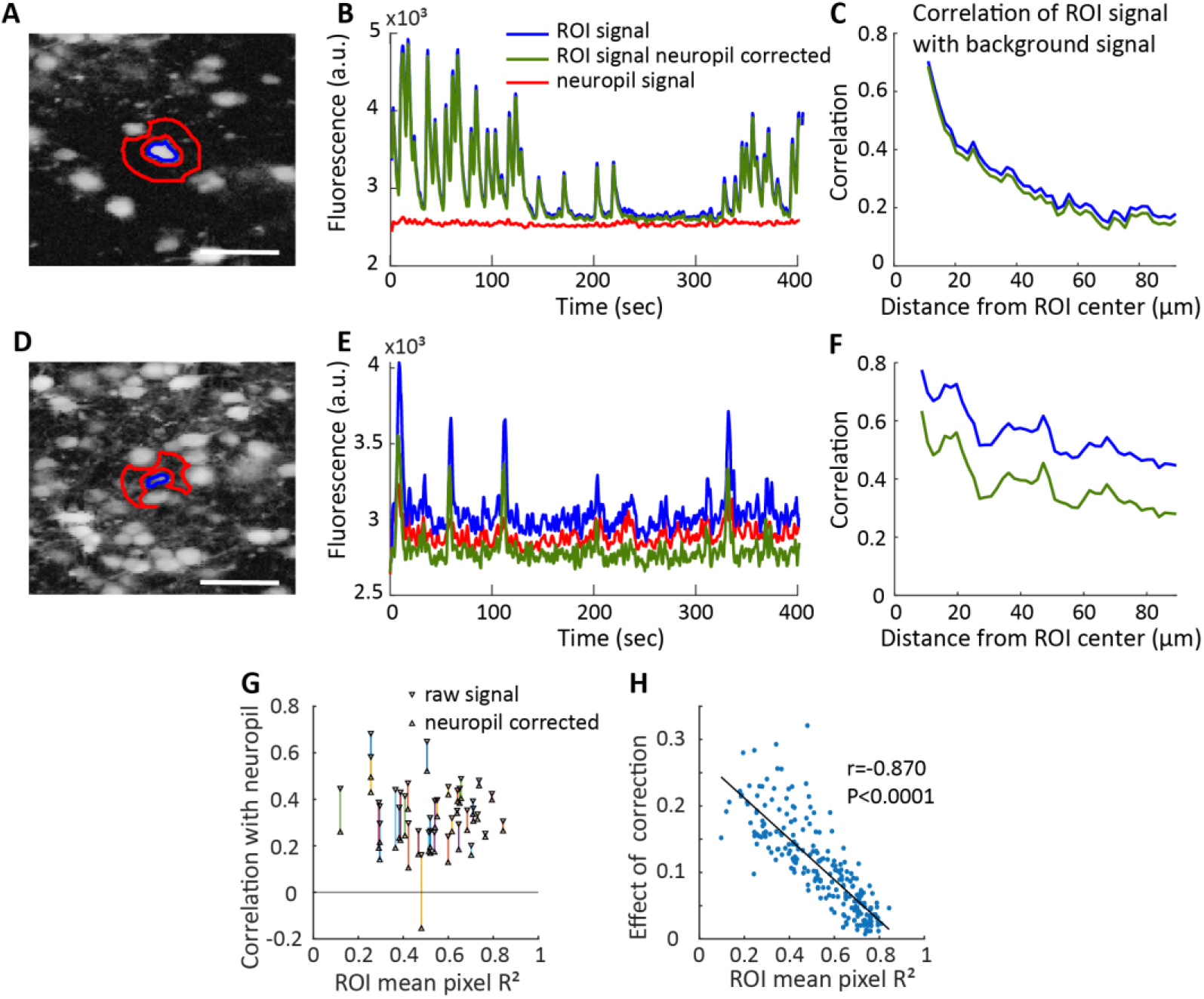
Neuropil estimation from surrounding pixels. **A**) Spectral image with an example ROI (blue contour) and surrounding background (red contour). **B**) Fluorescence signal of the example ROI of **A**. The neuropil signal is almost non-existent and neuropil correction does not change the ROIs’ signal much. **C**) Correlation of neuropil signal with ROI signal depends on the distance from the ROI. The fluorescence signal was extracted from rings at increasing distances around the example ROI, excluding other ROIs. Those background signals were correlated with both the raw signal of the ROI (blue line), and with the ROIs’ neuropil corrected signal (green line). Neuropil correction slightly decreases the correlation of the background signal. **D**) Spectral image with a different example ROI (blue contour) and surrounding background (red contour). **E**) In this ROI the neuropil signal caused more contamination than in the example of **A**. Neuropil signal was larger because more dendrites are present, and the ROI itself has a less strong signal. The neuropil correction seems necessary for this ROI. **F**) The neuropil correction decreases the correlation with the surrounding background signal. **G)** Correlation with neuropil signal of multiple ROIs, before and after neuropil correction. The distance between these shows the ‘neuropil correction effect’. **H)** The neuropil correction effect is significantly smaller when an ROI has a stronger mean pixel R^2^.

To test how our neuropil subtraction alters the signal, we correlated the pixel trace of ROIs with the pixel traces from the surrounding area (Fig. 9A-F). We found that in our data, neuropil correction barely changed the ROI signal in most cases (Fig. 9A-C). In some cases, however, the signal improved after neuropil subtraction (Fig. 9D-F). There may be multiple reasons why neuropil subtraction has less effect in our hands than in studies by others (Pachitariu et al., 2017). Experimental conditions such as the number of neurons that are labeled, the brain region that is imaged or the calcium indicator that is used will influence to what extent neuropil contaminates the signal (Pachitariu et al., 2017). An additional possibility is that when ROIs with a high mean pixel R^2^ are selected, those ROIs that have a weak signal and are strongly influenced by neuropil are excluded. To test this directly, we plotted the effect of neuropil correction on the signal extracted from ROIs against their mean R^2^ (Fig. 9G, H). This revealed that indeed, ROIs with higher mean pixel R^2^ are barely affected by neuropil correction while those with low mean pixel R^2^ are strongly affected. Thus, selection of ROIs with high mean pixel R^2^ has the added advantage that it reduces the need for neuropil subtraction.

### Automated retrieval of the same ROIs in chronically recorded datasets

One of the great strengths of multi-photon microscopy of calcium responses in neuronal cell populations is that changes in neuronal activity can be followed at the single cell level over prolonged periods of time. To achieve this, it is essential to identify the same ROIs in chronic recordings of the same brain region. In very large datasets containing many recordings, this is a daunting task to do by hand, even after automated ROI selection. We therefore developed a toolbox for automated matching of ROIs in chronically recorded datasets. It involves the registration of the sequential recordings with each other, through translation and rotation of the spectral images. Next, overlapping ROIs are searched for in all possible pairs of recordings in the series (i.e. each recording is compared to all other recordings). By thresholding for a minimal amount of overlap between each pair of ROIs in both directions, only ROIs with similar shapes and sizes are matched. Finally, merging all the matched pairs of ROIs results in a “matching matrix” containing the matched ROIs that are found in the series of recordings.

An example of ROI matching in three sequential recordings of V1 neurons, performed with 2-4 weeks in between sessions, is shown in Fig. 10. In these color-coded images, many cells can be retrieved in all three recordings. However, some neurons are not retrieved in one or two of the recordings, as indicated by the triangles. This may be caused by various factors, such as the precise angle and depth of the recording, changes in viral expression of the calcium indicator or the lack of activity of a neuron during one of the recordings.

**Figure 10.**
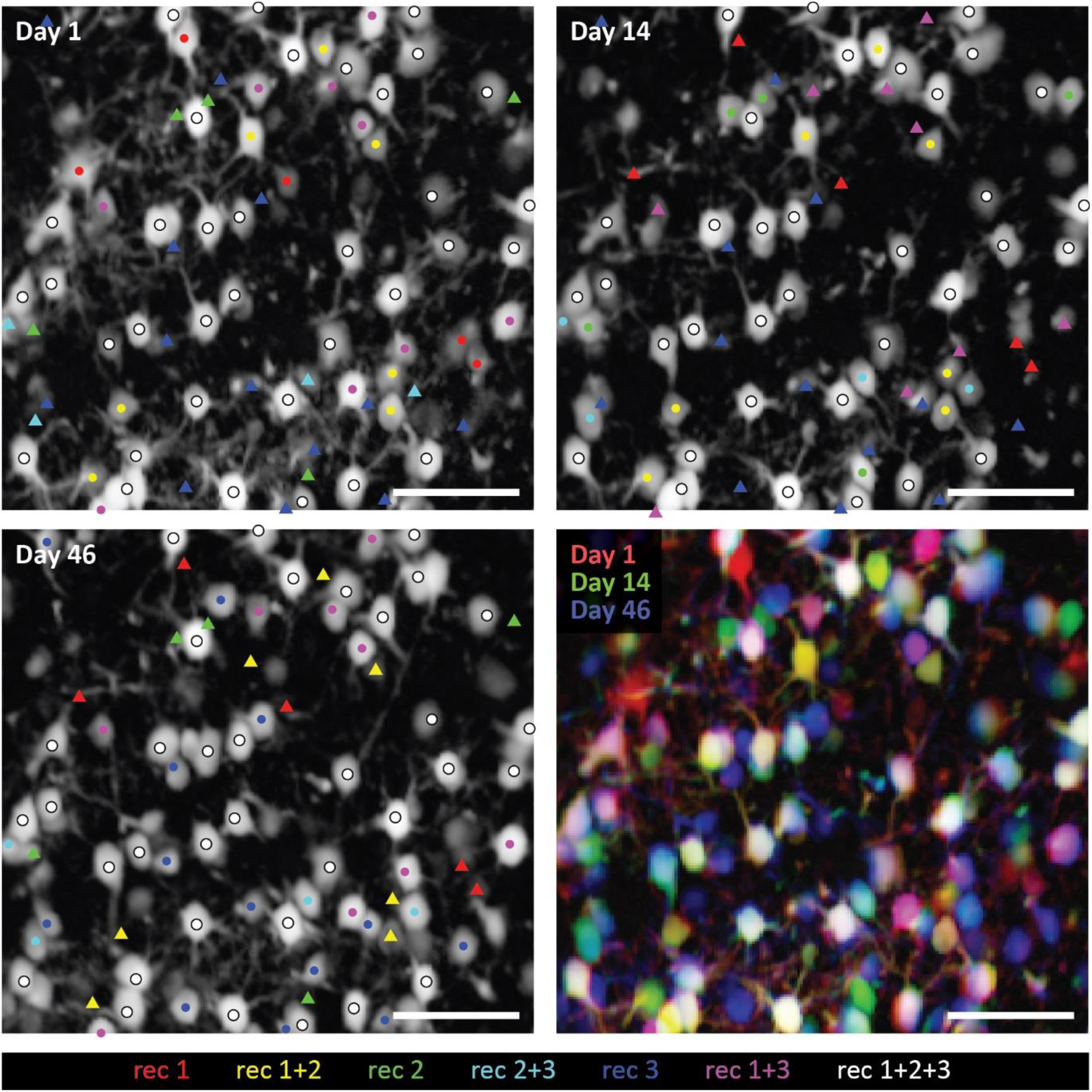
An example of matched ROIs in three sequential recordings. An ROI’s center of mass (CoM) is shown with a dot. The dot’s color shows in which recordings (‘rec’) it was retrieved. The triangles denote the position of some CoMs of ROIs that were present in other recordings, but missed in the recording in which the triangle is shown. The color denotes in which recording the ROI was present. The right bottom figure shows the three spectral images overlaid with red, green and blue colors, resulting in the same color mixing as the ROI CoM dots. Scale bar: 40µm.

### Comparing ROIs between recordings

For a quantitative assessment of the number of ROIs that can be matched between recording sessions we analyzed data from 5 mice that were imaged 7, 8, 12, 13 and 12 times respectively, over a period of 2-5 months. This resulted in 21, 28, 66, 78 and 66 possible pairs of recordings respectively, in which we investigated the number of matched ROIs. With all data of the five mice pooled together, 37.7% ± 21.3% of the ROIs were matched between recordings pairs, corresponding to 94± 57 ROIs. The percentage of matched ROIs was calculated as the percentage of matched ROIs in the recording with the least ROIs per recording-pair.

The number of matches was influenced by the time between two recordings. Fig. 11A shows that there was a significant correlation between the number of matched neurons and the time between recordings in two of the three mice (Spearman correlation, for mice 1 and 2 P<0.005, for mice 3, 4 and 5 P<0.0005, r=-0.6, −0.35, −0.66, −093, −0.68 for mice 1 to 5 respectively. The fits shown in the figure are exponential fits (y = *a* * e^*b* * x^) with the initial value (*a*) at 75, 71, 53, 56, 61(%), and decay rate (*b*) at −0.0084, −0.0036, −0.0171, −0.0160, −0.0123 for mice 1 to 5 respectively). This suggests that it is not only variations in the angle or exact location of the recordings that determines whether ROIs can be matched, but probably also slower, biological processes such as viral expression of the calcium indicator, learning- or age-induced changes in neural activity, cell death or anatomical changes of the brain. Indeed, Fig. 11B shows that correlations between the registered spectral images decreases over time (spearman correlation P=0.0006 for mice 1, for all other mice P<0.0005, r=-0.68, −0.58, −0.61, −0.93, −0.82 for mice 1 to 5 respectively), suggesting that the structure of the imaged location and activity patterns and GCaMP6f labeling of the neurons alter over a period of weeks to months. Changes in GCaMP6f labeling and/or neuronal activity is supported by the observation that more ROIs were detected on later imaging sessions for some mice (Spearman correlation P=0.2, P=0.008, P=0.51, P=0.97, P=0.015, and r=0.56, 0.83, −0.21, −0.01, 0.68 for mice 1 to 5 respectively) (Fig. 11D). Finally, we tested what fraction of ROIs could be tracked in multiple sessions. For pooling the data from the 5 imaged mice, we were constrained by the minimum number of recordings of the mice. We therefore limited the number of sessions to 7, choosing the sessions in such a way that the number of days between the first and last recording was close to 65. We then calculated in how many imaging sessions the ROIs from recording 1 were found back in subsequent sessions (Fig. 11C). If ROIs were found in 6 or less sessions, this was not necessarily in consecutive ones. As shown in Fig. 10, ROIs can be missing for some sessions, and then show up again. We found that 22% of all ROIs could be matched in all 7 recordings, averaged over the five mice, which corresponded to 213 neurons, or 213*7=1491 separate ROIs in total. 28% of ROIs (354 ROIs) could not be matched with ROIs from any other recording. Together, these results show that the developed pipeline is highly efficient in identifying ROIs and tracking them over many imaging sessions with little effort of the user.

**Figure 11.**
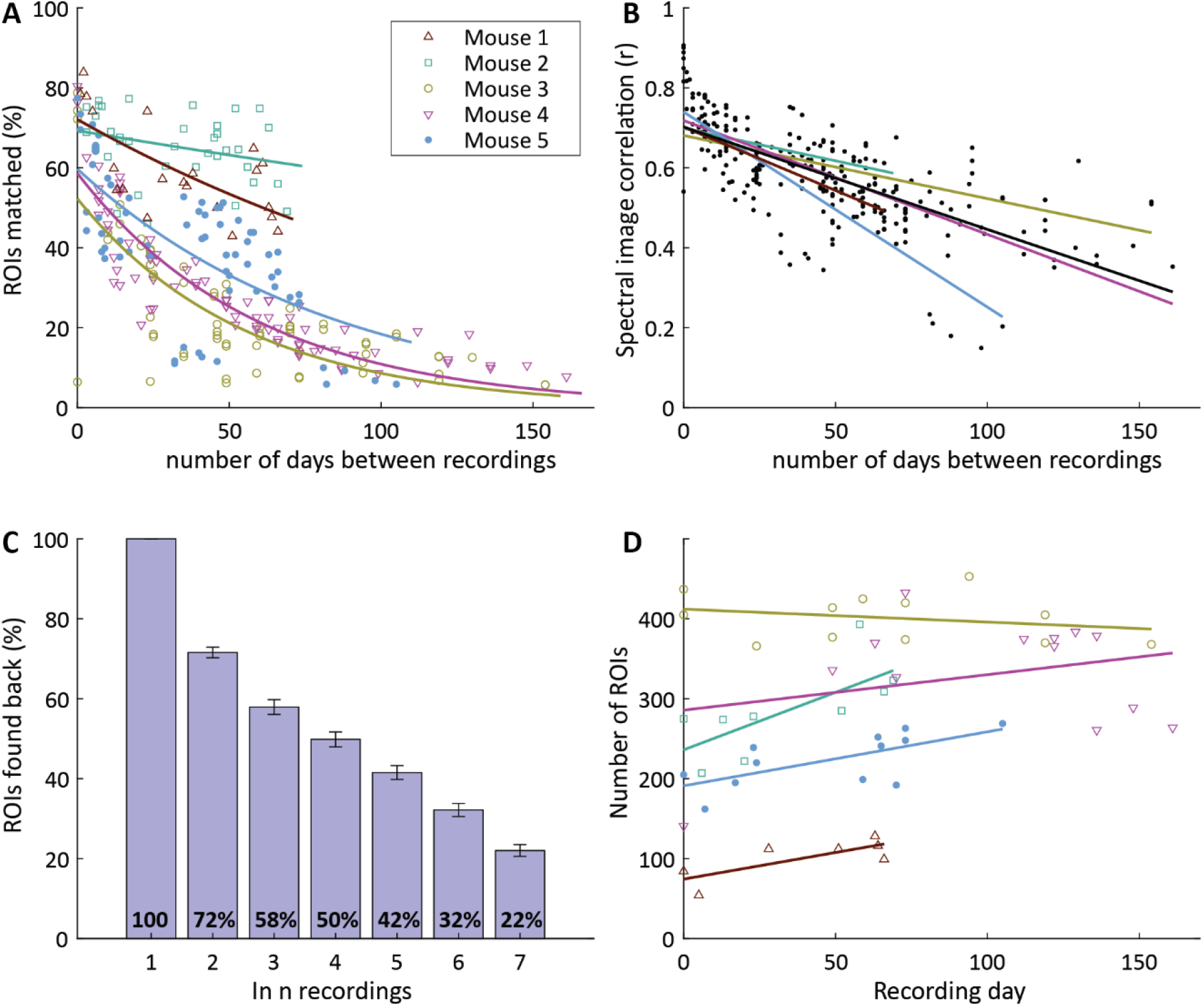
Chronic tracking of ROIs. **A)** The number of matches between recordings decreases if there is more time between the recording pairs. **B)** Spectral correlation for each recording pair. Recordings that are recorded closer together in time have better correlation coefficients. This shows that the imaged brain changes over time. **C)** Checking in how many recordings the ROIs from recording one were found back in all seven recordings. To be able to pool all the mice together seven recordings were analyzed per mouse. Recordings were chosen so that the time between the first and last recording was as close to 65 days as possible. The number of putative neurons in each condition were: 1146, 792, 606, 516, 421, 319 and 213 for conditions 1 to 7 respectively. Error bar shows standard error of the mean for the five mice. **D)** The number of ROIs found in the mice recordings did increase significantly for mouse 2 and 5 (Spearman correlation: P<0.05, for mouse 1, 3 and 4 the P values were 0.20, 0.51 and 0.97 respectively).

## Discussion

Here we describe SpecSeg, a novel open-source toolbox for automated ROI selection, ROI editing, neuropil correction, signal extraction and chronic matching of ROIs. SpecSeg ROI selection is based on clustering of co-active pixel traces combined with morphological filtering to identify somata and neurites alike. Pixel correlations have also been used in several other approaches for ROI selection (Kaifosh et al., 2014; Smith and Häusser, 2010; Spaen et al., 2019). In our implementation, however, the first step of ROI selection involves calculating the low (0.013-0.5Hz) frequency cross-spectral power components. These spectral components can be visualized, yielding exceptionally clear images of the local functional anatomy that include neuronal cell bodies and neurites. Underlying these low-frequency components are bouts of neuronal activity, which cause a prolonged rise in the fluorescence of the calcium indicator. In contrast to the high-frequency components, the low-frequency components are less sensitive to noise. Moreover, neuropil shows little low-frequency activity thus enabling the efficient separation of neuronal structures from the background. This first step of our ROI selection process requires only few initial constraints, permitting inclusion of ROIs with irregular shapes or sizes. During the second step, a threshold is set on the pixel correlations within the ROI, which leads to a robust separation of ROIs containing highly correlated pixels and rejection of areas of pixels in overlapping neural elements. Additional filters for morphological properties of the ROIs such as size or roundedness can be included. Adjustment of these constraints makes it possible to select ROIs around neuronal cell bodies, dendrites or axons. These constraints are intuitive and easy to use, making the SpecSeg ROI selection process very versatile and user friendly.

The pipeline also includes a graphical user interface, facilitating quality control of the selected ROIs. This interface comprises an alternative method for visualizing pixel correlations within the ROI, aimed at detecting whether multiple cells are present within one ROI. This makes it easy to split or reject ROIs if necessary. Also, ROIs can be added if the experimenter believes that cells are missed, and the quality of their signal can be assessed immediately.

We used the NeuroFinder datasets (http://neurofinder.codeneuro.org/) to compare the efficiency of the automated ROI selection procedure with other ROI segmentation packages available. This revealed that SpecSeg scored lower than other frequently used packages such as Suite2P (Pachitariu et al., 2017) and CaImAn (Giovannucci et al., 2019). This may have been caused by the settings we used for ROI selection, by limitations of our ROI selection approach, or by what are considered “ground-truth” ROIs in the Neurofinder datasets (or a combination of these factors). When we performed quality control on the Neurofinder ground-truth ROIs, we noticed that in a considerable subset the pixel traces within the ROI showed very little correlation. This implies that most of the signal extracted from pixels in such ROIs did not represent the signal from the ROI itself, but rather from surrounding neuropil. In contrast, SpecSeg only included ROIs with a high signal to noise (SNR) ratio. This also included ROIs that were too dark to be detected by assessing the average fluorescent signal alone and were missed in the Neurofinder ground-truth ROIs. This means that SpecSeg selects the most relevant ROIs but misses ROIs of neurons that are silent during the recording. A similar observation was made in a study comparing ROIs identified automatically using CaImAn and those identified by human annotators. It was found that matching ROIs were preferably those with a high SNR (Giovannucci et al., 2019).

To achieve a more refined set of ROIs that more closely match ROIs selected by human annotators, both CalmAn and Suite2P include a final step to exclude spurious ROIs by training classifiers (Giovannucci et al., 2019; Pachitariu et al., 2017). In SpecSeg, we chose to rely as little as possible on human annotation (directly, or indirectly by using trained classifiers) because our analyses revealed that this results in the inclusion of many ROIs with low SNR. We think it is debatable whether such ROIs should be included in the data analysis, as they may contaminate the actual neural code of the imaged neurons. When electrophysiological approaches are used, units with a signal that cannot be reliably separated from background noise would certainly not be included in the analysis. Only in some instances it is useful to know that a neuron is included in the dataset but does not show any activity, for example when chronically tracking the activity of individual neurons. In such cases, the option to manually include neurons using the SpecSeg graphical interface when they are not detected by the activity-based approach can solve the issue. An additional problem with using trained classifiers is that they will need to be retrained when other structures than neuronal cell bodies are to be identified for ROI selection, such as axons or dendrites.

We included a neuropil subtraction approach in our pipeline that works similarly to previously described methods (Chen et al., 2013; Khan et al., 2018; Tegtmeier et al., 2018). It calculates the signal from a donut-shaped area around the ROI, which is then subtracted from the signal derived from the ROI. We noticed that for most ROIs, neuropil correction did not significantly improve the neuronal signal. Of course, the level of neuropil contamination depends on many factors, including the brain region imaged, the strength of the neuronal signals, the density of labeling and the synchronicity of neuronal activity. After more thorough analysis, we noticed that ROIs with high mean pixel R^2^ were much less contaminated by the neuropil signal than those with low mean pixel R^2^. It should be noted that in ROIs with low mean pixel R^2^, neuropil subtraction may actually introduce an artefact in the signal, again illustrating the importance of removing such ROIs from the analysis.

As the use of miniscopes in freely behaving animals is rapidly increasing in labs around the world, we also optimized the pipeline for the analysis of calcium-imaging data obtained using this approach. We found that SpecSeg identified many ROIs overlapping with those detected by CNMF-E, and that the signals extracted from the ROIs defined by both methods were strongly correlated. In addition, different ROIs were detected by both approaches. However, many of these ROIs did actually show partial overlap and identified (parts of) the same neurons. This is probably caused by the fact that CNMF-E can create overlapping ROIs around neurons. The signal extraction procedure then separates these overlapping signals. In contrast, SpecSeg selects those parts of the neurons that do not overlap as ROIs and then extracts the signals from these ROIs. We therefore expect that in most cases, results using either method will be very similar, despite apparent differences in part of the selected ROIs. The main advantage of SpecSeg is its graphical user interface that provides extensive insight in the activity profiles of the ROIs and facilitates quality control and post-hoc ROI selection and editing.

It is difficult to compare the speed of the ROI selection process to other software packages available. However, we found that motion correction using NoRMCorre was the slowest step in the process, taking up approximately half the time of the total automated ROI selection process. This implies that the actual ROI selection process is not a bottleneck. Importantly, we found that the speed of ROI selection was more than sufficient for all practical purposes except on-line ROI selection.

One of the strengths of calcium imaging approaches is that one can follow the responses of individual neurons over prolonged periods of time. To facilitate this approach, we developed a tool for automated matching of ROIs in whole series of sequentially recorded calcium imaging sessions. Using this approach, about half of the ROIs between two randomly selected pairs of recordings could be matched. As already mentioned, one cause for not retrieving ROIs in all recordings is that some neurons are silent during some recordings, and thus not included by our activity-dependent ROI selection approach. Also, ROIs may be missed due to slight misalignments of the imaged brain region. Interestingly, we found that when the time between two recordings was longer, the number of matched ROIs was reduced. This shows that biological factors also play a role in the success rate by which neurons are detected in multiple recordings. Over time, the brain may slightly change its shape, causing some neurons to be excluded from the field of view. Moreover, neurons may die, or lose or gain expression of the viral vector. The most interesting reason seems to be that neurons may not be active in some recordings, in line with the finding that when mice perform the same task over prolonged periods of time, neuronal activity patterns reorganize over time resulting in the recruitment of different sets of neurons (Driscoll et al., 2017). Together, these issues may explain why studies in which the chronic tracking of activity of individual neurons are still sparse. Our automated matching approach will make this exciting possibility of multi-photon calcium imaging more accessible.

In conclusion, SpecSeg is a powerful, complete and open-source pipeline for ROI selection, signal extraction and chronic ROI matching that can be used on a variety of single- and multi-photon calcium imaging data. Its main advantages over several existing calcium imaging toolboxes are the ease of use and simplicity, the intuitive way ROIs are selected and constrained, the selection of ROIs that represent neurons whose responses can be well separated from background noise, the possibility to select ROIs of various shapes and sizes, its graphical interface for ROI editing and its use for analyzing both miniscope and multi-photon microscope calcium imaging data.

## Materials and methods

### Mouse experiments

To develop the ROI selection tools described in this study, we made use of two-photon imaging data from mice that were repeatedly imaged in as yet unpublished behavioral studies. Here we describe the methodology developed for the analysis of such data. All animal experiments were approved by the institutional animal care and use committees of the Royal Netherlands Academy of Arts and Sciences. We used male and female mice that were 2-7 months of age. The mice were C57Bl/6 mice, or offspring of Ai14 mice (Cre-dependent tdTomato reporter mice, strain 007908) crossed with mice expressing Cre in vasoactive intestinal polypeptide (VIP)-expressing or somatostatin (SOM)-expressing interneurons (Jackson Laboratories, www.jaxmice.jax.org, strains 010908 and 013044 respectively). All animals were kept in a 12 h reverse day/night cycle with access to food and water ad libitum. Experiments were carried out during the dark cycle.

### Viral injections for two-photon microscopy experiments

Mice were injected with a viral vector driving expression of the genetically encoded calcium sensor GCaMP6f in neurons (AAV2/9.syn.GCaMP6f, UPenn Vector Core facility). Anesthesia was induced with 5% isoflurane and maintained at 1.6% isoflurane in Oxygen (0.8L/min flow rate). Mice were administered Metacam (1mg/kg subcutaneously (s.c.), for analgesia) and dexamethasone (8 mg/kg s.c.) to prevent cerebral edema/inflammation and Cefotaxim (25 mg/kg s.c.) as antibiotic profylaxis, after induction of anesthesia. Mice were head-fixed on a stereotax, scalp and soft tissue overlying the visual cortex were incised and the skull exposed. A small hole was drilled in the skull overlying the center of primary visual cortex (V1). A pulled capillary with AAV2/9.syn.GCaMP6f was inserted vertically through this hole to a depth of 200-400 um from the brain surface. Approximately 20 to 100 nl of virus (titer ∼10E12 viral genomes per ml) was injected slowly using a Nanoject and the hole was covered with bone wax. During the surgery, the temperature was maintained with euthermic pads. Respiration was monitored to adjust depth of anesthesia. Eyes were protected from light and from drying using Cavasan eye ointment. Once the window was made the exposed dura was continuously kept moist with artificial aCSF, consisting of a solution of 125 NaCl, 10 Hepes, 5 KCl, 2 MgSO_4_, 2 CaCl_2_, and 10 Glucose, in mM. Later the scalp was sutured, and the animal let to recover from anesthesia.

### Handling and habituation

Once an animal recovered from the viral injection, and before window implantation, animals were handled daily for 5 minutes (or until they started to groom while being handheld) to reduce handling stress during later training. Next, animals were trained and habituated for 3 days with head-restrainment in the training setup with a running wheel. After habituation, animals were placed in a two-photon microscopy setup. Once the mice were comfortable with the setup, they performed a visual detection task.

### Cranial Window Surgery for two-photon microscopy

One month after viral vector injection, mice were anesthetized again as described above. Mice were head-fixed on a stereotax and scalp and soft tissue overlying the visual cortex were incised and the skull exposed. A metal ring (5mm inner diameter) was fixed on the skull centered on V1, with dental cement. A cranial window was made inside the ring and the dura was exposed. The cranial window was then covered with a double coverslip (to reduce brain movements under the microscope) and fixed to the metal ring using dental cement. Animals were allowed to recover after the dental cement dried. After a minimum of 2 weeks of recovery, mice were submitted to further handling and training. During the training and recording periods described below, animals were typically in the setup 5 days per week.

### Two-photon imaging

For imaging we used a Neurolabware standard microscope (CA, USA) equipped with a Ti-sapphire laser (Mai-Tai, Spectra-physics, CA, USA). A black cloth was used to cover the objective in order to prevent light coming from the monitor to the objective. Two-photon laser scanning microscopy was performed at 920 nm and neurons were imaged with 16x water-immersion objective (0.8NA) with computer-optimized optics of 1.6x magnification.

Two-photon calcium-imaging sequences were recorded in awake behaving mice and saved in a continuous binary format (sbx, Neurolabware). The dimensions of the images were 812 by 512 pixels, with 16 bit unsigned integer pixel depth. These files are associated with a metadata file (mat, MATLAB) that defines, among other parameters, pixel dimensions, number of channels, number of sections, and number of frames recorded. For two-photon GRIN lens imaging in the visual thalamus, we used a 4x objective (0.2NA) with computer-optimized optics with 3.2 magnification. All further processing was done with MATLAB (Mathworks™).

### Visual stimulation during two-photon microscopy

Stimuli were presented on a gamma-corrected Dell-P2314H 23” full HD LED monitor, placed 15 cm in front of the mouse. Stimuli were made with custom-made MATLAB scripts. Receptive field size of each region of interest (ROI) was estimated by reverse correlation after presenting 3 black and 3 white squares (7.5° -size degrees) simultaneously at pseudo random locations on the screen. The stimuli were repeated for 10-15 times at each location. The duration was 0.5 s, the interstimulus was an isoluminant gray screen, duration 1.5 s, contrast: 1.0 and maximum luminance was kept to 20% of screen max (max luminance). For visual stimulation, various stimuli were used, depending on the behavioral experiment.

### Gradient-index lens surgery

For single-photon miniscope experiments, animals were anaesthetized with isoflurane (1.5% in O_2_ and air) and placed in a stereotaxic apparatus (David Kopf Instruments) on a heated pad (37°C), injected with Metacam (1mg/kg subcutaneously) as analgesic, dexamethasone (8 mg/kg subcutaneously) to prevent inflammation, and saline (80 ml/kg subcutaneously) to prevent dehydration. A craniotomy and durotomy were made above the region of interest. To image calcium dynamics in premotor cortex, a total of 500nl AAVdj.CaMKIIa.GCaMP6s (Titer max. 10E12 genomes per ml, Stanford Neuroscience Gene Vector and Virus Core) was injected at four locations 100µm off center of premotor cortex (AP:2.3 ML:0.35 DV: −0.3) using a Hamilton-syringe (100nl/min). A GRIN objective lens (1.8mm diameter, Edmund Optics Ltd.) was placed on top of the brain. To image striatum, a custom-built robotized stereotaxic arm was used to slowly lower (300nm/min) a 25G needle to ease lens implantation and reduce tissue damage above the ventral striatum (AP:1.1 ML:1.1 DV:-4.5). A total of 500nl AAVdj.hSyn.GCaMP6s was injected at two different location 100µm off center of ventral striatum (AP:1.1 ML:1.1 DV:-4.5), followed by lowering (100nm/min) of a GRIN relay lens (0.6mm diameter, Inscopix Inc.). The gap between lens and skull was covered with cyanoacrylate glue and secured to the skull using Superbond dental cement (Sun Medical Company Ltd.). Using dental acrylic cement, a head bar was placed further caudal and skin was glued to the cement headcap. The lens was protected by a layer of Twinsil Speed (Picodent GmbH) silicon.

For two-photon imaging in the visual thalamus, animals were anesthetized as described above and injected with a total of 100nl AAVdj-hSyn.GCaMP6s (Titer ∼10E12 genomes per ml, Stanford Neuroscience Gene Vector and Virus Core) at two locations (AP:2.2 & 2.4, ML:2.1) and three depths per location (DV:2.35, 10nl; 2.5; 20nl, 2.7; 20nl). A doublet GRIN lens (1mm diameter, 0.5mm working distance, GRINTECH) was implanted right above the visual thalamus as described above for striatum, centered in between the two injection locations (AP:2.3, ML:2.1, DV:2.1) and fixed on the skull using Superbond dental cement. A custom-made head restriction ring was placed around the GRIN lens and fixed on the skull using Superbond and Tetric EvoFlow^®^ (A1). The GRIN lens was protected by a custom-made 3D-printed cap that could be screwed on the head restriction ring.

### Baseplating and miniscope imaging

Four to eight weeks after virus injection and GRIN lens placement, mice were head restricted using a custom-built device (that included a running belt to allow for locomotion) and fluorescent signal was assessed. The miniscope was lowered above the implanted GRIN lens and, once individual cells were detectable in the field of view, a baseplate was secured to the cement head cap using dental acrylic cement (coated with black nail polish). Recording imaging sessions were performed in an open-field arena (30cm x 30cm). Prior to each imaging session, mice were head restricted, and the miniscope was attached to the baseplate. Calcium dynamics were recorded with a frame-rate acquisition of 15Hz, and data was stored in videos files with a resolution of 752 (width) x 480 (height) pixels (AVI format).

### Miniscope recording and preprocessing

Raw AVI files were spatially down sampled by a factor four (reducing frames to 376 x 240 pixels) and stored as TIFF files. Rigid and non-rigid motion in imaging data was corrected using NoRMCorre (Pnevmatikakis and Giovannucci, 2017), followed by neuron and signal extraction using either SpecSeg or CNMF-E (Zhou et al., 2018). Every cell included in the analyses was verified by visual inspection. CNMF-E output was manually cleaned using a custom-written GUI in MATLAB (2019b). Components with non-circular contours, spatial contours smaller than neuron size, or artifacts in temporal traces were excluded. Duplicate copies of individual cells were identified by overlapping contours and temporal activity was subsequently merged.

### SpecSeg data analysis code

The code for all data analysis procedures described below is available at Github: (https://github.com/Leveltlab/SpectralSegmentation). A flowchart of the analysis pipeline is shown in Fig. 1 and is also present in the Github repository.

### ROI selection based on cross-spectral power

We developed a pipeline for ROI extraction that works as follows: First, the images are cropped to remove border artifacts before being aligned using rigid registration with NoRMCorre (https://github.com/flatironinstitute/NoRMCorre), a toolbox provided by the Simons Foundation (Pnevmatikakis and Giovannucci, 2017). We adapted the entry function of this toolbox in order for it to work with files in .sbx format, produced by the Neurolabware microscope that we used, and integrated it in the pipeline. After visual inspection to ascertain that the images are well aligned, the image sequences are transposed to place time in the first dimension and width * height in the second dimension (StackTranspose.m). This makes the processing of pixel traces much more efficient in the time domain. This data is saved in binary format for later use with “_Trans.dat” as extension. In addition, we also save a downsampled version of this file with “_DecTrans.dat” as extension. Downsampling is perfomed with “decimate” (MATLAB) to a sampling rate of ±1Hz. Next, the cross-spectral power of the fluorescent signal between neighboring pixels over time is calculated. To achieve this, the data is first detrended and for each half-overlapping time window (60 seconds) the data is convolved with a hamming window. Cross-spectral density functions are calculated from the discrete Fourier transforms of these pixel-trace segments for each pixel with its eight neighbors and averaged over all time windows. Additionally, the total variance is calculated for each pixel from their average auto-spectral density function. Then, the cross-spectral power functions are normalized (see formula 1) with their respective variances.

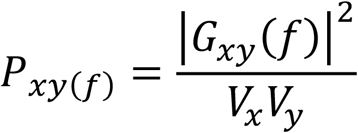

***Formula 1***. *Cross-spectral power: Where P*_*xy*_ *(f) is the average normalized cross-spectral power, G*_*xy*_ *(f) is the cross spectral density between x and y for frequency component f, and Vx and Vy the average variance of x and y respectively*.

Finally, the normalized cross-spectral power functions of a pixel with its eight neighbors are averaged. The result is a 2D matrix representing cross-spectral power at each frequency component for all pixels. We used this to generate an image for each spectral component. This data is saved in a separate file with “_SPSIG.mat” as extension.

Cross-spectral images are generated for spectral components between 0.017 Hz to 0.5 Hz, from image sequences recorded at a sampling rate between 10 and 30 Hz (Fig. 2). Pixels within active neurons display strong cross-spectral power below 0.4 Hz whereas background neuropil is usually non correlated making neurons clearly visible in these images (Figs. 2, 3). The images for cross-spectral components between 0.017 and 0.4 Hz are therefore used to extract ROIs (spectral.m). For each cross-spectral image in the selected frequency range, a series of morphological constraints are applied to find as many non-overlapping ROIs as possible (getSpectrois.m). First, all maxima (cross-spectral peaks) in the image are detected and sorted according to magnitude. A fraction (0.25-0.4) of these peaks with the highest values are selected and sorted in decreasing order. Based on this selection, a square area of pixels (Voxel; 50×50 pixels) is sampled for each peak, centered on the peak and contours are detected using contourc (MATLAB) (Fig 3D). Closed contours containing the selected peak are selected and constrained by a set of criteria:

1. To avoid that multiple peaks from the same neuron are selected, the minimal distance between peaks in each ROI should be greater than a defined threshold (20 pixels).
2. To ensure that the magnitude of the peak is well above background level, the peak should be greater than (95%) of the pixel range within a voxel.
3. Given the expected size range of cell bodies, the minimum and maximum area (number of pixels) should be between 40-400 pixels (a pixel is ±1.5µm^2^ within our field of view: ±1mm).
4. Roundedness; The relationship between the area of a contour and the length of its circumference (4*Pi* Area / Circumference^2) should be greater than 0.6 (1 = completely round, 0 = any shape), These parameters can be adapted depending on the type of ROI that a user is interested in and the magnification of the images. To select cell bodies, for example, the roundedness should be greater than 0.6 and the number of pixels should be below 100 at low magnification. Because the criteria are simple and self-evident, optimal values can easily be established with some experimentation. Depending on the density of active neurons and their processes in an image sequence, contours around correlated pixels may still represent overlapping or adjacent neural elements. Therefore, the ROIs are further restrained based on the assumption that pixels in adjacent cell bodies are not highly correlated. To achieve this, all pixel traces within the original ROI are correlated with the averaged down-sampled trace of eight neighboring pixels at the original local maximum. Based on this calculation, the ROI is constrained to an area containing pixels with correlations larger than the half maximum of the total range (Fig. 3E). In principle, this is a computationally expensive approach, but it is feasible because the numbers of pixel traces in a preselected ROI are limited in number and the traces are decimated to 1Hz. For analysis and selection purposes we also saved the mean pixel R^2^ of an ROI. To calculate this we averaged the squared pixel correlations within the constrained ROI.

### Miniscope background correction

We included a background correction tool (BackgroundSubtractSbx.m) for miniscope calcium imaging data, which is applied after the motion correction step. A unique background estimation image is created for every frame by filtering a padded version of that frame. This background estimation is subtracted from the frame. Before subtraction, the values of the background estimation are lowered by a fixed amount to prevent underexposure of the data in the final result. There are three different filter kernels to choose from: a Gaussian, a disk, or a donut. A disk of 111×111 pixels (∼200um) was chosen in the example data (Fig. 4E-F, Fig. 6). The gap in the donut filter prevents the background estimation from containing signal of a neuron at the position of the filter. We found, however, that the difference in results compared to a circular filter was very small. A Gaussian filter provides very similar results to a smaller disk filter, and much of the miniscope data is already out of focus in a way that resembles a Gaussian kernel. This is why the circular kernel is preferred over the Gaussian. The size of the filter can be edited by the user, based on the data’s scale. When selecting the donut-shaped filter, both the gap size and outer radius of the filter can be adjusted. Background subtraction takes more processing time to calculate with larger filters or larger datasets. To diminish processing time, the background can be calculated using a downscaled version of each frame. An optional feature in the background subtraction process is data smoothing. To decrease sensor banding noise and increase correlation between neighboring pixels, data can be smoothed with a Gaussian in either the vertical, horizontal or both directions. When the correlation between neighboring pixels is increased, it becomes easier to detect ROIs with low mean pixel R^2^. This smoothing was not used in the example data. All filtering parameters can first be tested on a small subset of the data with the BackgroundSubtractSbxExample.m script.

### User interface for ROI refinement

To give users control over the selected ROIs, we developed a graphical user interface to reject ROIs outside a preferred range of properties (RoiManagerGUI.m). Additionally, the user interface allows the user to manually delete or keep ROIs. In rare cases, some ROIs have areas with pixels that have low or negative correlations. This may indicate that the ROI contains signals from multiple neuronal sources, or that registration was suboptimal and the neuron was not located at the ROI during the entire recording. We therefore developed the option to split these ROIs (Fig. 7B-E). The user interface creates four reference points at the distal edges of the ROI. The fluorescence signal from these reference points is then correlated with the signal of each pixel in the ROI. This creates four correlation values per pixel of the ROI. These are used to create a color-coded image in the user interface, indicating the location of different signal sources in the ROI. The pixels in the ROI can then be subjected to k-means clustering. After setting the number of clusters the user can decide to split such an ROI in two or more ROIs or to delete part of the ROI. The user can also add more reference points if clustering needs to be improved.

We also included the option to create new ROIs in the user interface, for example when neurons were missed by the automated analysis. These new ROIs can be added in two different ways. One option is to manually draw the contour of the ROI. The other option is to let the computer draw the ROI, based on the cross-spectral image (or any other user-defined background image, such as the fluorescent image) and a threshold set by the user (Fig. 7A).

### Signal extraction and neuropil subtraction

To extract the calcium signals for each ROI (retrievesignals.m), all pixel traces within the ROI contour are averaged. The signals are neuropil corrected by subtracting the averaged signal of a neuropil area from the ROI signal. A donut-shaped neuropil area is created for each ROI. To achieve this, a small buffer area with a width of 2 pixels (∼2 µm) is first added around each ROI. The neuropil area is defined by first enlarging the ROI by 20 pixels (∼20 µm) using a circular filter, after which the central and adjacent ROIs and their buffer areas are excluded from this area. To prevent overcorrection, the neuropil signal is multiplied by 0.7, in accordance with previous studies (Chen et al., 2013; Khan et al., 2018; Tegtmeier et al., 2018).

### Spike estimation

The extracted ROI signal can be converted into an estimate of a spike train (DeconvolveSignals.m) with the MLspike toolbox (Deneux et al., 2016), which has to be downloaded via their github link (https://github.com/MLspike). The spike train is then saved in the same format as the regular calcium signals.

### Identification of the same ROIs in chronically recorded datasets

Finally, we include a toolbox to identify the same ROIs in chronic recordings of the same brain region (ChronicMatching.m). It first registers the spectral images of the recordings to ensure that each ROI will be located at exactly the same position in all recordings. To achieve this, the spectral images are normalized so that all values are in the range from −1 to 1. Next, the registration of the spectral images is done by repeated translation and rotation. Each recording is registered to a reference recording chosen by the user. Preferentially, one of the middle recordings is used as a reference in order to minimize the difference between the reference and the recordings to be registered. The distance over which recordings need to be translated is calculated by doing a 2D cross-correlation between the reference image and the spectral images of the other recordings. The cross-correlation is calculated with xcorr2_fft (Masullo, 2020), which is much faster than the MATLAB built-in cross-correlation function. After the translations are applied, the images are padded with zeros to maintain the same dimensions.

To correct the rotation, each recording is rotated using bicubic interpolation over a range of different angles, from 1° clockwise to 1° counterclockwise in steps of 0.05°. Each rotation is compared to the reference image with a 2D correlation and the best fit is applied if the correlation is at least 0.005 higher than the original image. Several rounds of translation and rotation are applied until no significant further improvement is achieved.

After registration, ROIs between all chronic recordings within an experiment are matched. A minimum percentage of overlap threshold is set. In the example presented here, we chose 67.5% based on the best balance between false positives and false negatives as determined by inspection with the chronic viewer user interface. The ROI matching is represented in a 2D matrix (the “match matrix”), consisting of a column for each recording and rows with the ROI numbers in each recording that are matched to each other. If no match for a particular ROI is found in a recording, the cell is kept blank.

To create the match matrix, the following three steps are taken:

1. For each ROI in each recording, the overlapping ROIs from all other recordings are identified. The percentage of the reference ROI that is covered by each overlapping ROI of other recordings was calculated (“overlap”) and saved as a putative match if it is above the overlap threshold.
2. Next, the overlap for each pair of putatively matched ROIs is calculated in the opposite direction (i.e. the percentage of overlapping ROI that is covered by the reference ROI). The average of the two overlap values are averaged, and putative matches whose average overlap is below the overlap threshold (in our example 67.5%) are discarded. Calculating the overlap in these two simple steps automatically takes many parameters of the neurons into account: if the sizes, shapes or positions of two ROIs differ significantly the overlap value will always be small.
3. To obtain the final matching matrix, the data needs to be merged in order to match the same ROIs in all recordings, not just in pairs of recordings. The merging process includes ‘mutual linking’: if an ROI from recording A is matched only to recording B, but that ROI from recording B has matches to A and C, it is implied that the ROI from recording A is also matched to the ROI in recording C. This mutual linking increases the number of matched ROIs.

The results can be checked in a user interface (ChronicViewer.m). The matches can be verified by selecting a cell in the match matrix, which will light up the ROI contours of the match. Clicking on ROIs in the main image will show the ROI numbers and match information in a table. The match matrix can be edited if necessary.

## Acknowledgements

The authors thank Emma Ruimschotel for technical assistance, and the Mechatronics department and Animal facilities of the NIN for their services. This project has received funding from the European Union’s Horizon 2020 Research and Innovation Programme under Grant Agreements No. 785907 (HBP SGA2) and 945539 (HBP SGA3). We thank Dr. Karl Deisseroth for providing us with AAVdj.CaMKIIa.GCaMP6s and AAVdj.hSyn.GCaMP6s viral vectors.

## Competing interests

The authors report no competing interests.

## Notes

https://github.com/Leveltlab/SpectralSegmentation

### Competing Interest Statement

The authors have declared no competing interest.

### Summary of Updates

The toolbox has been extended so that it works really well with single-photon miniscope imaging data. This is shown in a new figure (Figure 6) comparing SpecSeg with CNMF-E. In addition, the software has been updated to improve installation on Apple Mac computers.

